# Non-photopic and photopic visual cycles differentially regulate immediate, early and late-phases of cone photoreceptor-mediated vision

**DOI:** 10.1101/2020.02.15.950915

**Authors:** Rebecca Ward, Joanna J. Kaylor, Diego F. Cobice, Dionissia A. Pepe, Eoghan M. McGarrigle, Susan E. Brockerhoff, James B. Hurley, Gabriel H. Travis, Breandán N. Kennedy

## Abstract

Cone photoreceptors in the retina enable vision over a wide range of light intensities. However, the processes enabling cone vision in bright light (*i.e.* photopic vision) are not adequately understood. Chromophore regeneration of cone photopigments may require the retinal pigment epithelium (RPE) and/or retinal Müller glia. In the RPE, isomerization of all-*trans*-retinyl esters (atRE) to 11-*cis*-retinol (11cROL) is mediated by the retinoid isomerohydrolase Rpe65. An alternative retinoid isomerase, dihydroceramide desaturase-1 (DES1), is expressed in RPE and Müller cells. The retinol-isomerase activities of Rpe65 and Des1 are inhibited by emixustat and fenretinide, respectively. Here, we tested the effects of these visual cycle inhibitors on immediate, early and late phases of cone photopic vision. In zebrafish larvae raised under cyclic light conditions, fenretinide impaired late cone photopic vision, whereas emixustat-treated zebrafish unexpectedly had normal vision. In contrast, emixustat-treated larvae raised under extensive dark-adaption displayed significantly attenuated immediate photopic vision concomitantly with significantly reduced 11-*cis*-retinaldehyde (11cRAL). Following 30 minutes of light, early photopic vision recovered, despite 11cRAL levels remaining significantly reduced. Defects in immediate cone photopic vision were rescued in emixustat- or fenretinide-treated larvae following exogenous 9-*cis*-retinaldehyde (9cRAL) supplementation. Genetic knockout of *degs1* or retinaldehyde-binding protein 1b (*rlbp1b)* revealed that neither are required for photopic vision in zebrafish. Our findings define the molecular and temporal requirements of the non-photopic and photopic visual cycles for mediating vision in bright light.

## Introduction

Photopic vision is the physiological function mediating vision in well-lit conditions. Daytime vision requires cone photoreceptors to remain functional in the presence of light. This capability requires regeneration of cone photo-pigments in the retina prior to and during light onset. However, our understanding of the biological process contributing to immediate, early and late phases of cone photopic vision is inadequate.

### Visual Cycles in RPE and Müller cells

Sustained vision in vertebrates depends upon a supply of 11-*cis*-retinaldehyde (11cRAL) chromophore to regenerate photobleached visual pigments. The canonical visual cycle is the enzymatic pathway in the retinal pigment epithelium (RPE) that recycles visual chromophore (1). The retinoid isomerase in RPE cells is RPE65, which converts all-*trans* retinyl esters (atREs), synthesized by lecithin-retinol acyltransferase (LRAT), to 11-*cis*-retinol (11cROL) (2–4). Null mutations in the mouse *Rpe65* or *Lrat* gene cause the virtual absence of 11cRAL within the neural retina (5,6). *Rpe65*^*−/−*^ mice exhibit massive accumulation of atREs in the RPE, while retinoids are almost undetectable in RPE of *Lrat*^*−/−*^ mice (7,8). Notably, there is still uncertainty as to if, and when, the RPE65 visual cycle contributes to cone photopic vision.

Under daylight conditions, the estimated rate of 11cRAL synthesis by RPE cells is much slower than the rate of chromophore consumption by rods and cones (9). Despite the non-contribution of rods to daylight vision, rhodopsin continues to photoisomerize in bright-light exposed retinas. Cones must therefore compete with rods for chromophore when demand is highest. As a possible adaptation, cones can regenerate cone-opsin pigments by uptake of either 11cRAL or 11cROL, while rods can only uptake 11cRAL (10–12). Müller glial cells of the neural retina are the likely source of this 11cROL (13). Dihydroceramide desaturase (DES1; encoded by *DEGS1*) and multifunctional *O*-acyltransferase (MFAT) represent another retinol isomerase—retinyl-ester synthase pair, present in Müller cells, which may mediate 11cROL synthesis. The ‘*isomerosynthase’* activity of DES1—MFAT is present in retinas from cone-dominant chickens and ground squirrels, but undetectable in retinas from rod-dominant mice and cattle (9,14). This activity correlated with the presence of 11-*cis*-retinyl-esters (11cREs) in the neural retina, which are undetectable in mice and cattle retinas. Interestingly, synthesis of 11cROL by DES1 is increased through interactions with cellular retinaldehyde binding protein (CRALBP; encoded by *RLBP1*) an 11cRAL/11cROL carrier protein expressed in the RPE and Müller glia (15).

In contrast to the evidence supporting a role for DES1 in cone chromophore regeneration, a recent study reported that recovery of cone electrophysiological sensitivity, in isolated retinas following a photobleach, was similar in *Des1*^*+/+*^ and Müller-cell conditional *Des1*^*−/−*^ mutant mice (16). This suggests Müller DES1 plays no role in the generation of retinoids to recover mouse cone electroretinograms. However, uncertainty remains regarding the overall contribution of DES1 to the cone visual cycle as significant DES1 activity remained in retinal lysates of these Müller-cell *Des1*^*−/−*^ mice.

Additionally, the cone electrophysiological studies were conducted in rod-dominant mice whereas DES1 was identified in Müller cells from cone-dominant chicken. Thus, uncertainly remains as to whether species-specific retinoid isomerase pathways exist to meet cone chromophore demands.

A third mechanism postulated to replenish cones with chromophore is the photopic visual cycle, a process that regenerates 11cRAL by a light-dependent mechanism. The non-visual opsin, retinal G protein-coupled receptor (RGR) opsin is present in RPE and Müller cells. Previous reports show RGR-opsin is integral to light-dependent biochemical formation of 11cRAL that reconstitutes rhodopsin visual pigment in rods (17). RGR-opsin appears essential for the recovery of cone electrophysiological sensitivity as retinas from wild-type mice under sustained background illumination maintained cone sensitivity, while *Rgr*^*−/−*^ mice exhibited diminishing cone sensitivity during light exposure (18). Notably, all these studies assess cone electroretinography (ERG) and not cone vision. It is well recognized that electrophysiology does not absolutely correlate to functional vision. For example, patients receiving gene therapy for RPE65 defects show no improvement in ERG but show significant improved ability to navigate an obstacle course (19).

### In Vivo Cone Models

Most visual cycle knowledge was ascertained by studying retinae of rod dominant mice. Consequently, investigations of the visual cycles enabling cone vision are less advanced. Zebrafish larvae provide novel opportunities to study cone photopic vision and the supporting visual cycle processes (20) due to easy genetic and pharmacological manipulation, and cone-dominant vision until 15 dpf, the stage when rods become functional (21,22). Previous studies investigating visual cycle components in zebrafish relied on transient morpholino knockdown. Zebrafish *rpe65a* knockdown significantly reduced, but did not completely attenuate 11cRAL synthesis, supporting the presence of an alternate pathway for chromophore synthesis in zebrafish retinae (20). Knockdown of either Cralbpa or Cralbpb in zebrafish resulted in reduced visual behavior (23) and reduced light sensitivity (24).

### Visual Cycle Modulation

Pharmacological modulators that inhibit or complement the visual cycles are potential treatments for retinal disease (25,26). Following photoisomerization, released atRAL must be delivered to RPE/Müller cells to regenerate 11cRAL to restore light sensitivity. Impaired atRAL clearance from photoreceptors leads to accumulation of bis-retinoid *N*-retinylidene-*N*-retinylethanolamine (A2E), a toxic by-product of atRAL and a pathological hallmark of age-related macular degeneration (AMD) and Stargardt disease (27). Emixustat (ACU-4429) competitively inhibits Rpe65 and hinders retinaldehyde-mediated destruction of photoreceptors by acting as a retinaldehyde scavenger (26). Inhibition of chromophore synthesis gives rise to unliganded ‘noisy’ opsins that stimulate the transduction pathway in the absence of light, greatly decreasing photoreceptor sensitivity (28). In *Rpe65*^*−/−*^ mice, chronic activation of signal transduction may cause photoreceptor degeneration (29). Free opsin can be targeted by chromophore-replacement therapy with 9cRAL, which bypasses the visual cycle defect to form isorhodopsin or iso-cone-opsins. Following 9cRAL treatment, light sensitivity is restored long-term in *Rpe65*^*−/−*^ mice with attenuated accumulation of atREs (30). Fenretinide (4-HPR), a derivative of retinoic acid is reported to inhibit DES1without affecting RPE65 activity (31–33). Fenretinide also competes with retinol for binding to retinol-binding protein (RBP4) in blood, causing a mild vitamin A deficiency (34). A1120 is a non-retinoid RBP4 antagonist which in mice reduces serum retinol levels in a dose-dependent manner (35) and reduces lipofuscin accumulation in *Abca4*^*−/−*^ mice through reduction of serum RBP4 and visual retinoids (34). Unlike fenretinide, A1120 does not act as a retinoic acid receptor alpha agonist. A1120 does not affect RPE65 isomerohydrolase activity in *ex vivo* experiments (34).

Previous visual cycle inhibition studies focused on measuring retinal function (*e.g.* electrophysiology) as the primary endpoint. Reduced cone photosensitivity was observed in live mice and isolated retinas treated with visual cycle inhibitors (32). To our knowledge, no previous studies investigated the relationship between visual cycle modulation and cone photopic vision. In the current study, we used visual behavior assays, visual cycle inhibitors and genetic approaches in zebrafish to understand the mechanisms regulating immediate (at light onset following dark adaptation, 0 minutes), early (after 30 minutes light adaptation) or late (after ~4-6 hours light adaptation) phases of cone photopic vision. Original findings were uncovered in relation to cone photopic vision, including*: i)* late photopic vision is significantly impaired by fenretinide, but not by emixustat, *ii)* immediate photopic vision is reliant on Rpe65, *iii)* 30 mins of light is sufficient to restore early photopic vision and surmount the RPE65 inhibitor emixustat *iv*) the RPE65-independent recovery of early photopic vision occurs with low levels of 11cRAL, *v*) exogenous 9cRAL is sufficient to restore photopic vision and *vi)* photopic vision does not rely on *degs1* or RPE-expressed *rlbp1b*.

## Results

### Fenretinide, but not emixustat, impairs late cone photopic vision in zebrafish larvae

To investigate the effects of visual cycle inhibitors on cone vision, we assayed the effects of pharmacological agents, previously used in human clinical trials (36,37), on the vision of 5 dpf zebrafish larvae. Previous studies demonstrate 5 dpf zebrafish rely exclusively on cones for vision (38). Emixustat is a potent RPE65 inhibitor; while targets of fenretinide include DES1 (the putative isomerase II) (32) and retinol-binding protein 4 (Rbp4) in blood (34). Larvae were raised under normal cyclic light conditions (14 h light, 10 h dark) for the duration of the experiment and treated with emixustat or fenretinide from 3 dpf. Drug was replenished at 4 dpf, and OKR analysis completed at 5 dpf between ZT 2.5 – ZT 6.5 **(Figure 1A)**. Surprisingly, late photopic vision was unaffected in 5 dpf larvae treated with up to 50 μM emixustat (24.3 ± 4.9 saccades per minute) **(Figure 1B).** In contrast, larvae treated with 10 μM fenretinide displayed significantly fewer (P<0.001) saccades per minute (7.7 ± 5), in comparison to vehicle controls (24.2 ± 8.2) **(Figure 1B).** Wholemount images reveal that 5 dpf larvae treated with emixustat or fenretinide fail to inflate their swim bladder, but otherwise have normal gross morphology **(Figure 1C). Figure 1D** depicts potential targets of emixustat and fenretinide in zebrafish. Zebrafish have three *rpe65* isoforms, two of which are expressed in the eye (20,39). Emixustat may inhibit RPE-expressed *rpe65a* or Müller-cell expressed *rpe65c* **(Figure 1D).** Fenretinide may inhibit DES1 isomerase activity in Müller cells/RPE, or inhibit uptake of atROL from the serum by blocking Rbp4-Transthyretin (TTR) interactions. In summary, in cyclic lighting conditions a broad-spectrum visual cycle inhibitor impairs late photopic vision, whereas a selective inhibitor of the canonical visual cycle did not affect vision.

**Figure 1:**
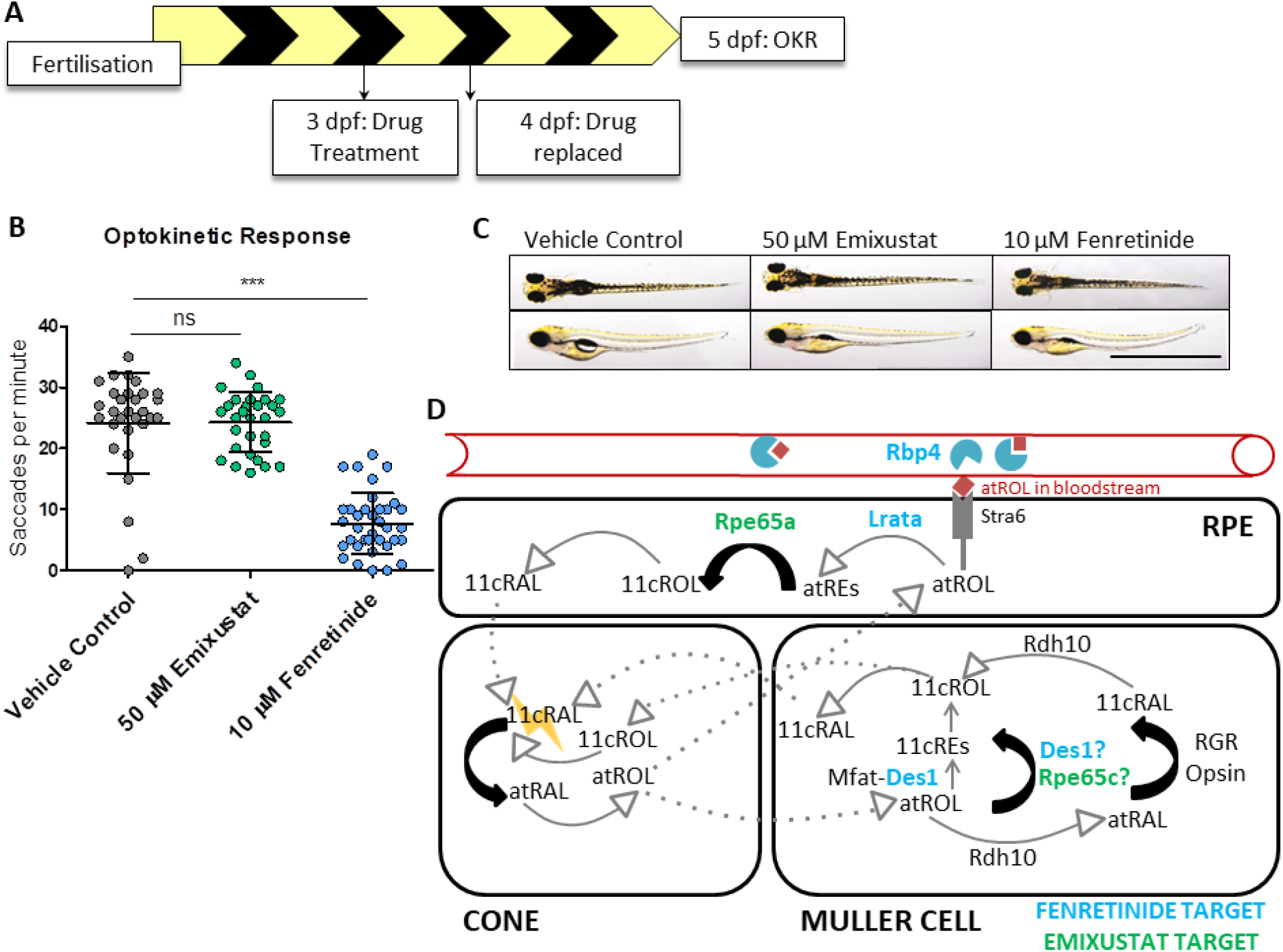
Fenretinide, but not emixustat, impairs late cone photopic vision in zebrafish larvae. **A)** Schematic representation of experimental workflow. Zebrafish larvae are treated initially at 3 dpf and drug is replaced at 4 dpf. Larvae are incubated under standard lighting conditions (14-hour light, 10-hour dark) until OKR analysis at 5 dpf. **B)** OKR in 5 dpf larvae following treatment with 50 μM emixustat and 10 μM fenretinide. Data were analyzed by a one-way ANOVA and Dunnett’s multiple comparisons post-hoc test where ns=not significant (p>0.05) and ***=p<0.001. N=30 larvae with 3 independent biological replicates. **C)** Dorsal and lateral brightfield microscopy images of larvae following treatment with 50 μM emixustat or 10 μM fenretinide at 5 dpf. Scale bar=2 mm. **D)** Schematic overview mapping all possible mechanisms of cone chromophore regeneration in zebrafish including the confirmed molecular targets of both emixustat and fenretinide. Proteins colored in blue depict fenretinide targets (Rbp4, Lrata & Des1) and those in green highlight (potential) targets of emixustat (Rpe65a, Rpe65c) in zebrafish.

### In dark adapted zebrafish, emixustat blocks immediate photopic vision and fenretinide exerts an additive effect

As Rpe65 is important for chromophore regeneration under dark conditions, we tested the visual cycle inhibitors in dark-adapted zebrafish larvae wherein larvae were raised under normal cyclic light conditions until 3 dpf, then treated with drug under constant dark conditions from 3-5 dpf. Visual behavior analysis was performed at 5 dpf between ZT 2.5 and ZT 6.5, wherein larvae were individually removed from wells and immediately subjected to OKR following a two-day dark adaptation **(Figure 2A).** Dose-response and dose-frequency studies revealed that 50 μM emixustat added at day 3 and replaced at day 4 caused the most significant reduction in OKR saccade number/minute (7.1 ± 4.6, p<0.001) compared to vehicle controls (20.9 ± 5) **(Figure 2B).** Under dark conditions, larvae treated with 10 μM fenretinide displayed a ~6-fold reduction in the ability to track 11 cm thick rotating stripes, whereas larvae treated with 50 μM emixustat had a ~2.5-fold decrease in OKR. Interestingly, when both isomerase inhibitors were combined, immediate photopic vision was completely abolished (1.2 saccades/minute, p<0.001) and reduced by ~22 fold, compared to vehicle controls (26.8 saccades per minute). **(Figure 2C).** Larvae treated with emixustat and fenretinide in the dark were morphologically normal, other than they did not inflate their swim bladder **(Figure 2D)** as also observed under a normal light/dark cycle **(Figure 1C).** An uninflated swim bladder is commonly observed in visually compromised zebrafish larvae (40,41). Interestingly, neither 50 μM emixustat nor 10 μM fenretinide induced obvious changes in retinal histology, particularly the photoreceptor outer segment structure and RPE, compared to vehicle controls **(Figure 2E-G, 2E’–2G’).** Furthermore, double cones stained with zpr-1 appear unaffected by fenretinide or emixustat treatment **(Figure 2E’’–2G’’).** Thus, emixustat impaired immediate photopic vision following dark adaptation **(Figure 2A and C)** but not late photopic vision **(Figure 1B)**, while fenretinide inhibited both immediate and late photopic vision.

**Figure 2:**
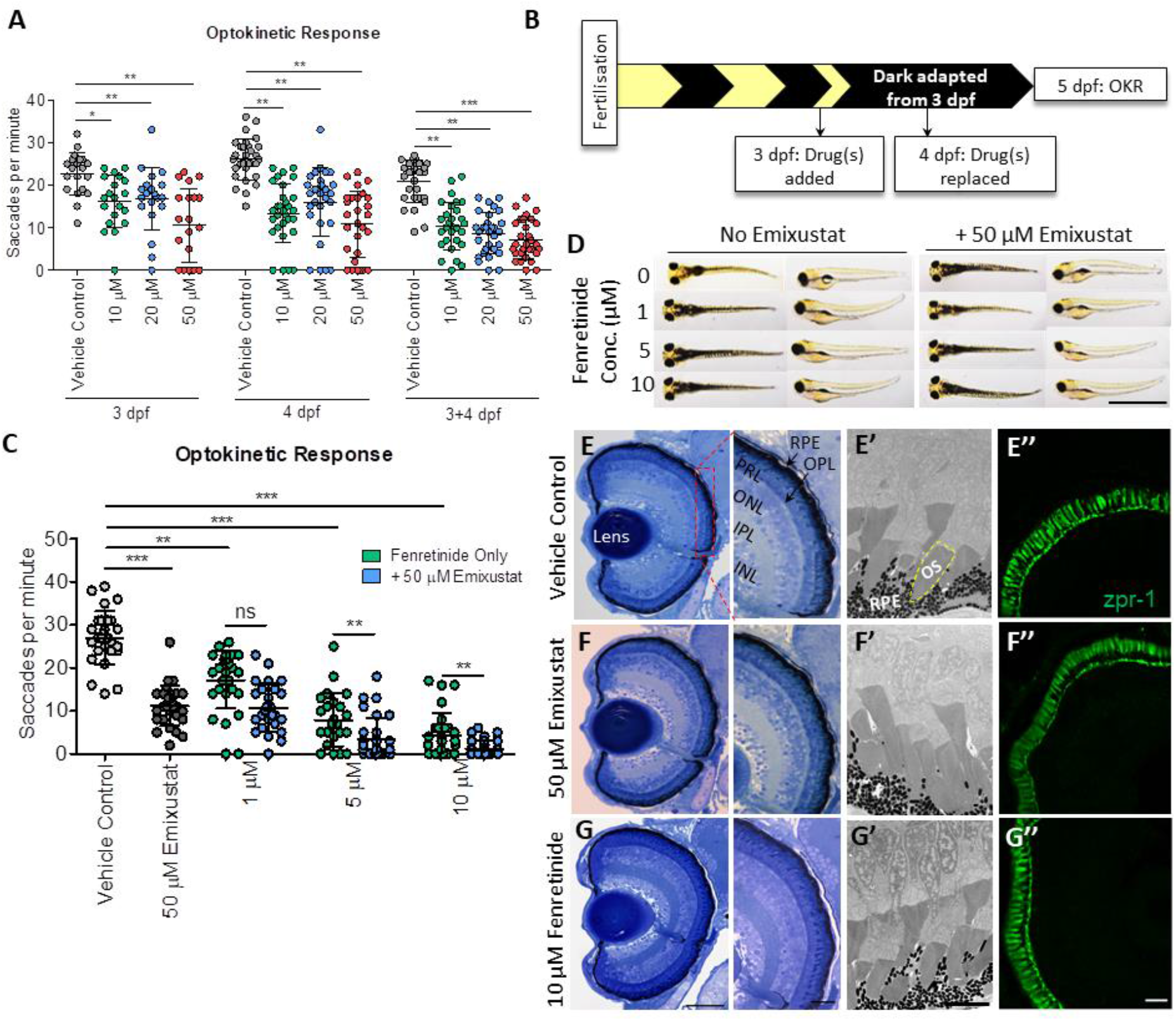
In dark adapted zebrafish, emixustat blocks immediate photopic vision and fenretinide exerts an additive effect. **A)** Optimization of dose and treatment time points for emixustat. Larval OKR at 5 dpf following treatment with 10, 20 or 50 μM at 3 dpf, 4 dpf, 3 and 4 dpf. Data were analyzed by two-way ANOVA and Bonferroni post-hoc test where *= P<;0.05, ** = p<;0.01 and ***=P<;0.001. N=30 larvae with 3 independent biological replicates. **B)** Schematic representation of experimental workflow. Zebrafish larvae are initially treated with emixustat and/or fenretinide at 3 dpf and drug(s) are replaced at 4 dpf. Larvae are incubated under dark conditions until OKR analysis at 5 dpf. **C)** Larval OKR at 5 dpf following combination treatment with 50 μM emixustat and/or 10 μM fenretinide at day 3 and day 4. Data were analyzed by two-way ANOVA and Bonferroni post-hoc test where ns=not significant and ** = p<; 0.01. **D)** Dorsal and lateral brightfield microscopy images of larvae at 5 dpf following treatment with 50 μM emixustat and/or 10 μM fenretinide. Scale bar = 2 mm. **E)** Retinal morphology of 5 dpf larvae treated with, 0.5% DMSO, 50 μM emixustat or 10 μM fenretinide. Retinal brightfield images taken with a X60 and X100 objective. RPE=Retinal Pigment Epithelium; PRL=Photoreceptor layer; OPL=Outer Plexiform Layer; ONL=Outer Nuclear layer; IPL= Inner Plexiform Layer; INL=Inner Nuclear Layer. Scale bar=50 μm & 20 μm, respectively. **E’ – G’)** High resolution images of photoreceptor outer segments (OS, yellow dotted line) and RPE in 5 dpf larvae. Scale bar = 5 μm. **E’’-G’’)** Immunohistochemistry on transverse section across lens at 5 dpf. Double cones were stained with zpr-1 antibody. Scale bar=20 μm.

### Visual cycle inhibitors modulate the profile of key retinoids in zebrafish

To assess the pharmacological specificity of the phenotypes in zebrafish, we measured the content of visual retinoids in zebrafish larval heads following pharmacological visual-cycle inhibition. Animals genetically or pharmacologically devoid of functional visual cycles can present with significantly altered levels of retinoids in the eye and liver (3,20,24,26,42,43). In agreement with the impaired immediate photopic vision we observe in zebrafish, we observed a significant reduction in 11cRAL, 11cROL and the 11cRE, 11-*cis*-retinyl palmitate (11cRP), following emixustat and fenretinide treatment, alone or in combination (p<0.001) **(Figure 3B).** Rpe65 uses atREs as its substrate, therefore, elevated levels could be expected in the presence of emixustat (2). Interestingly, atROL (23.2 ± 3.8 pmole/mg, p<0.01) and were significantly reduced in fenretinide-only treated larvae compared to the vehicle control (70.5 ± 27.3 and 20.8 ± 2.3 pmole/mg, respectively). These data suggest that emixustat and fenretinide impair immediate photopic vision following dark adaption in zebrafish by reducing 11cRAL levels.

**Figure 3:**
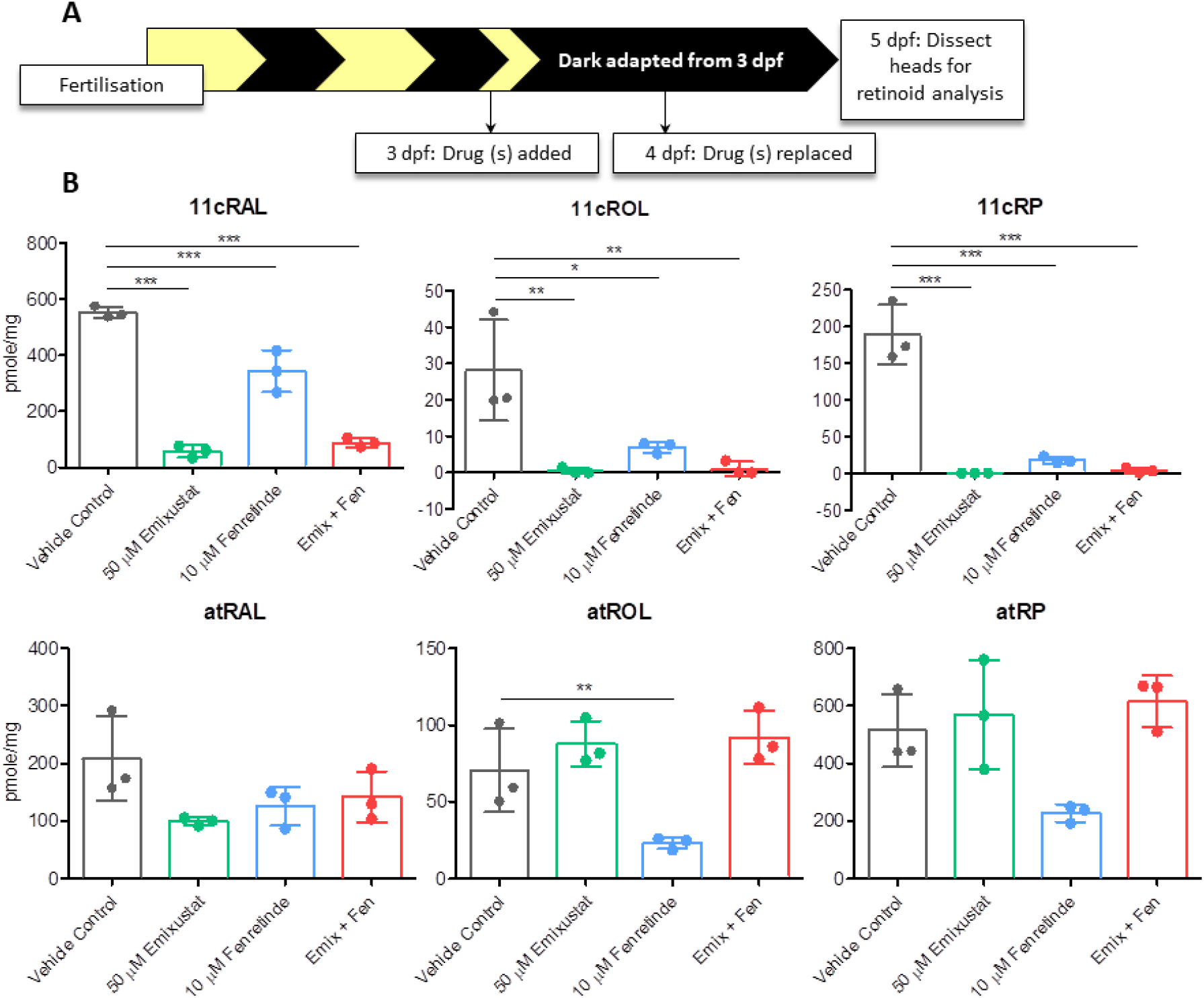
Visual cycle inhibitors modulate the profile of key retinoids in zebrafish. **A)** Schematic representation of experimental workflow. Zebrafish larvae are initially treated with 50 μM emixustat and/or 10 μM fenretinide at 3 dpf and drug is replaced at 4 dpf. Larvae are incubated under dark conditions until collection of larval heads for retinoid analysis at 5 dpf. **B)** Retinoid profiles of 5 dpf larvae treated with emixustat and fenretinide, alone and in combination. Bars represent the mean±SD of three independent experiments for each condition with 105 larval heads per biological replicate. Data were analyzed by a one-way ANOVA and Dunnett’s multiple comparisons post-hoc test where *=p<0.05, **=p<0.01 and ***=p<0.001.

### Supplementation with exogenous 9cRAL significantly improves immediate photopic vision in emixustat and fenretinide-treated larvae

As cone function is partially restored in *Rpe65*^*−/−*^ mice following treatment 9cRAL, a functional analogue of 11cRAL (44), we measured cone vision following treatment with exogenous 9cRAL. Larvae were treated with emixustat and/or fenretinide under dark-adapted conditions as described above **(Figure 2A).** In addition, larvae were treated with 9cRAL four hours following treatment with emixustat and/or fenretinide at both 3 and 4 dpf **(Figure 4A).** Immediate photopic vision in emixustat-treated larvae supplemented with 9cRAL improved significantly (16.1 ± 6.7 saccades per min, p<0.001), compared to emixustat-treatment alone (10.5 ± 4.8 saccades per min) **(Figure 4B).** We also observed approximately two-fold (6.5 ± 5.5 saccades per min, p<0.01) improved cone vision in fenretinide treated larvae following supplementation with 9cRAL **(Figure 4B).** 9cRAL supplementation improved vision in emixustat plus fenretinide-treated larvae approximately five-fold compared to emixustat and fenretinide-treated larvae without 9cRAL supplementation (5.3 vs 4.2 saccade per minute, respectively) **(Figure 4B).** No gross morphological changes were observed in larvae treated with 9cRAL in combination with fenretinide alone or fenretinide plus emixustat **(Figure 4C).** Interestingly, larvae treated with emixustat and 9cRAL inflated their swim bladders **(Figure 4C),** a characteristic not observed in larvae treated with emixustat alone **(Figure 2D).** Finally, treatment of larvae with exogenous 9cRAL altered retinoid profiles at 5 dpf **(Figure 4D)**. All 9-*cis* retinoids (9cRP, 9cROL and 9cRAL) and atRP significantly increased in emixustat and 9cRAL-treated larvae, compared to emixustat alone. Presence of small amounts of 9cRAL in samples not treated with exogenous chromophore is most likely a result of thermal isomerization **(Supplementary Figure 1).** Interestingly, no significant increase was observed in 11cRAL profile following 9cRAL supplementation. Thus, the reduction in immediate photopic vision in emixustat and fenretinide-treated larvae results not from toxic effects, but rather from visual cycle inhibition.

**Figure 4:**
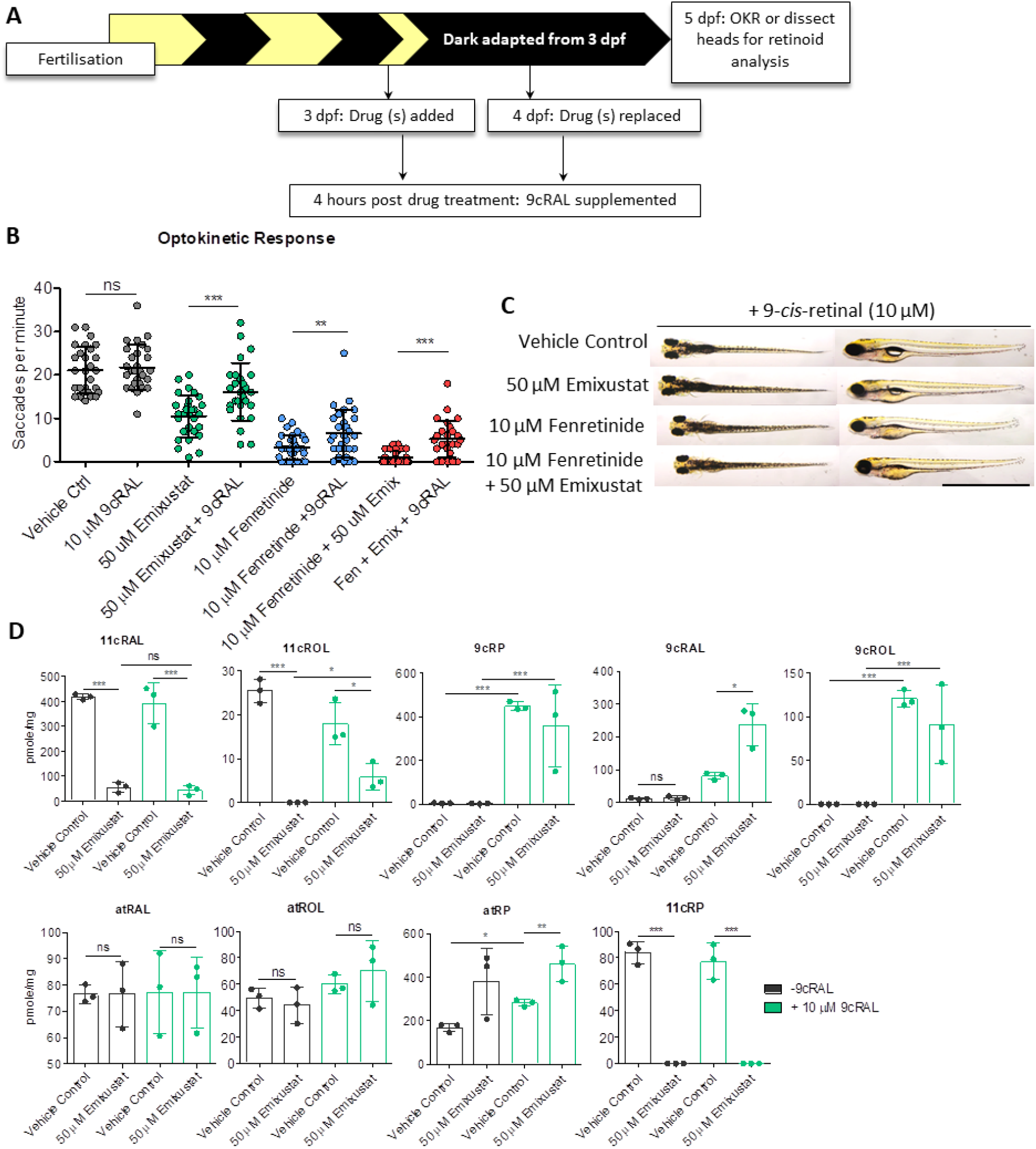
Supplementation with exogenous 9cRAL significantly improves immediate photopic vision in emixustat and fenretinide-treated larvae. **A)** Schematic representation of experimental workflow. Zebrafish larvae are treated initially at 3 dpf and the drug is replaced at 4 dpf. 9cRAL is added to the medium four hours following 50 μM emixustat and/or 10 μM fenretinide treatment on both 3 dpf and 4 dpf. Larvae are incubated in the dark until OKR analysis at 5 dpf. **B)** Larval OKR following addition of 10 μM 9cRAL to larvae prior treated with 50 μM emixustat and 10 μM fenretinide, alone or in combination at 3 dpf and 4 dpf. Data were analyzed by unpaired, two-tailed t-tests where *=p<0.05, **=p<0.01, ***=p<0.001. ns=not significant. N=30 larvae with 3 independent biological replicates. **C)** Dorsal and lateral brightfield microscopy images of larvae at 5 dpf larvae treated with 50 μM emixustat and/or 10 μM fenretinide in combination with 10 μM 9cRAL. Scale bar=2 mm. **D)** Retinoid profiles following exogenous 9cRAL supplementation in vehicle control and emixustat-treated zebrafish larvae. Bars represent the mean±SD of three independent experiments for each condition with 105 larval heads per biological replicate. Data were analyzed by unpaired, two-tailed t-tests where *=p<0.05, **=p<0;.01, ***=p<0.001. ns=not significant.

### Early photopic vision recovers in emixustat-treated larvae following exposure to light

Recent studies proposed the non-canonical visual cycle to rely on a Rpe65-independent isomerization event which relies on RGR-opsin and occurs during sustained exposure to visible light (18). We hypothesized that exposure of larvae to light for 30 minutes following a prolonged dark adaptation may activate the alternative pathway and modulate the impaired visual behavior in larvae treated with emixustat and fenretinide under dark-adapted conditions **(Figure 2C).** At 5 dpf, between ZT 2.5 – ZT 6.5, larvae were subjected to our standard OKR assay immediately after dark adaptation for two days (0 minutes) or following 30 minutes subsequent exposure to ambient light **(Figure 5A).** Pre-treatment with 50 μM emixustat partially suppressed immediate photopic vision in dark-adapted larvae (11.4 ± 7.6 saccades per minute, p<0.001) compared to vehicle controls (17.3 saccades per minute) **(Figure 5B).** In contrast, pre-treatment with 10 μM fenretinide virtually abolished immediate cone photopic vision in dark-adapted larvae (1.7 ± 3 saccades per minute, p<0.0.001) compared to the vehicle control (17.3 ± 4.5 saccades per minute) **(Figure 5B)**. To help discriminate if fenretinide acts via Rbp4 or Des1, A1120, a potent Rbp4 inhibitor was tested. Similar to fenretinide, A1120 suppressed immediate cone photopic vision (10.2 ± 6.6 saccades per minute). Subsequent exposure of untreated larvae to 30 minutes of light had minimum effect on early cone visual function versus untreated, dark-adapted larvae **(Figure 5B)**.

**Figure 5:**
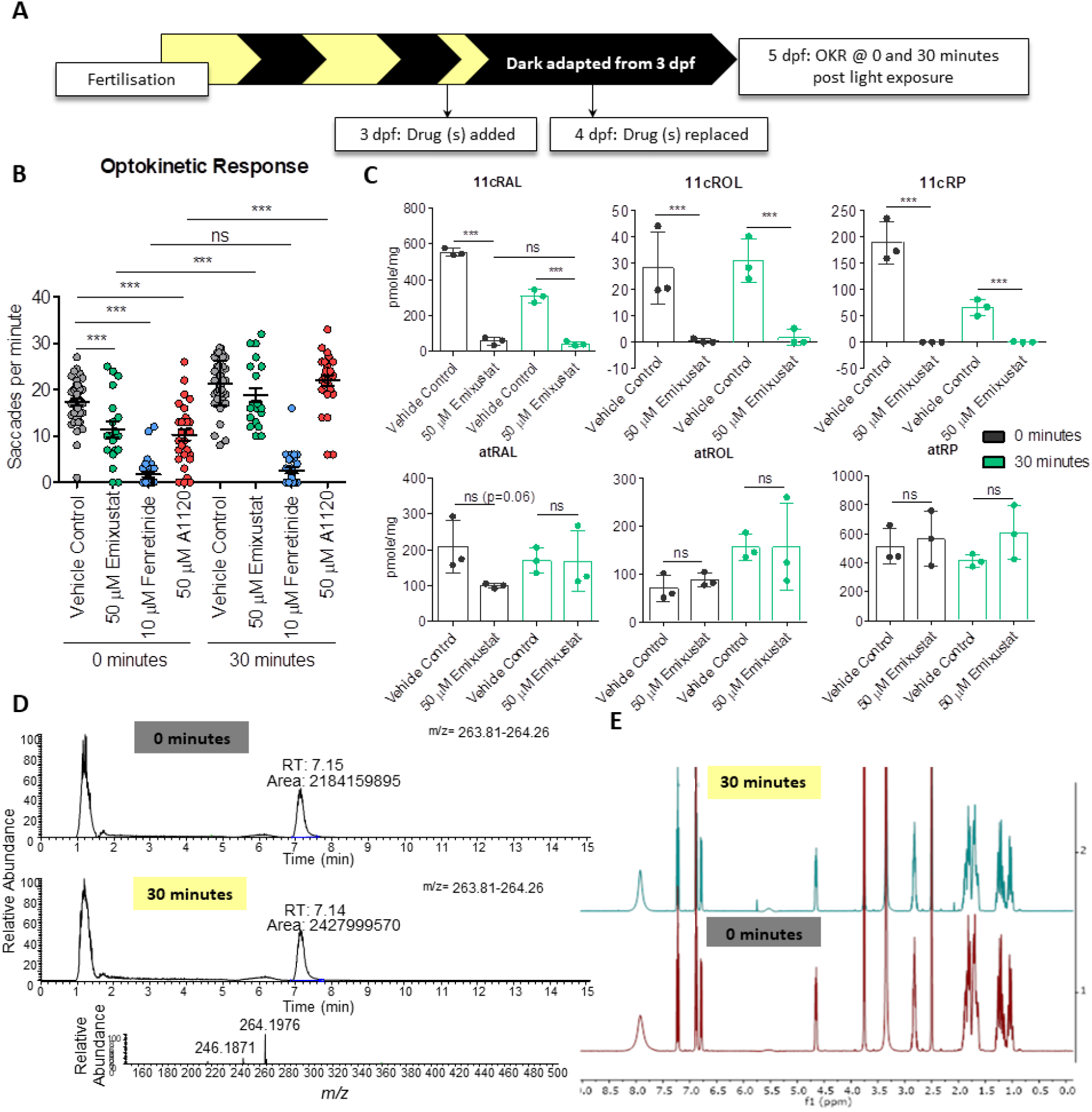
Early photopic vision recovers in emixustat-treated larvae following exposure to light. **A)** Schematic representation of experimental workflow. Zebrafish larvae are treated initially at 3 dpf and drug is replaced at 4 dpf. Larvae are incubated under dark conditions until analysis at 5 dpf. At 5 dpf larvae are subjected to OKR immediately following dark adaptation (0 minutes) or following 30 minutes of light (30 minutes). **B)** OKR of 5 dpf larvae treated with 50 μM emixustat, 50 μM A1120 or 10 μM fenretinide 0 and 30 minutes following light exposure. Data were analyzed by unpaired, two-tailed t-tests where ns=not significant, **=p<0.01 and ***=p<0.001. N=30 larvae with 3 independent biological replicates. **C)** Retinoid profiles of 5 dpf larvae treated with 50 μM emixustat following no light exposure (0 minutes) or 30 minutes light exposure (30 minutes). Bars represent the mean±SD of three independent experiments for each condition with 105 larval heads per biological replicate. Data were analyzed by unpaired, two-tailed t-tests where ns=not significant, **=p<0.01 and ***=p<0.001. **D)** Emixustat extracted ion chromatograms (at *m/z* 264, RT 7.1 min) and high resolution mass spectrum (*m/z* 264.1976 [M+H]^+^ highlighting no change in the mass spectrometry profile of emixustat exposed to 30 minutes of light and samples which received no light (0 minutes). **E)** NMR light sensitivity analysis of emixustat. Emixustat was dissolved in DMSO-d^6^ (600 μl) covering with aluminum foil until ^1^H NMR and HSQC experiments were performed (0 minutes). The NMR tube was left exposed to visible light for 30 minutes, followed by a second NMR analysis. (30 minutes). Peaks at 2.09 ppm and 5.76 ppm correspond to acetone and dichloromethane, respectively.

Interestingly, early photopic vision in emixustat-treated larvae was indistinguishable from vehicle-only treated larvae after 30-minutes exposure to light **(Figure 5B)**, despite lower 11cRAL levels in larvae treated with emixustat **(Figure 5C)**. Pre-treatment of larvae with fenretinide virtually abolished cone vision following 30 minutes of light exposure, similar to fenretinide-treated, dark-adapted larvae **(Figure 5B)**. Surprisingly, early photopic vision in A1120-treated larvae recovered to vehicle control levels (21 ± 5 saccades per minute) indicating that the block in early photopic vision observed following fenretinide treatment is not a result of inhibiting Rbp4 **(Figure 5B).**

The relatively normal cone OKRs in emixustat-treated larvae following light exposure raised the question if this correlated with increased 11cRAL chromophore levels. Retinoid levels were profiled in emixustat-treated larval heads at 0 and 30 minutes of light exposure **(Figure 5C).** Although vision recovered following 30 minutes light exposure **(Figure 5B)**, 11cRAL levels surprisingly did not change. Interestingly, atRAL levels returned to that of the vehicle control following 30 minutes indicating photoisomerization of light sensitive chromophore and activation of the phototransduction cascade.

As early photopic vision, in the presence of emixustat, recovered following exposure to ambient light for 30 minutes, there was a possibility of light-induced degradation or inactivation of emixustat. To test this possibility, we performed mass spectrometry analysis of emixustat before and after light exposure. No change in the relative abundance of the ion corresponding to emixustat (*m/z* = 263.81— 264.26) was observed **(Figure 5D)**. NMR analysis also demonstrated no chemical change to emixustat upon exposure to visible light. **(Figure 5E)**.

To confirm that early photopic vision was not mediated by reactivation of RPE65, larvae were treated in the dark with emixustat from 3 dpf and emixustat was removed at 5 dpf under dim red light, 30 minutes before OKR analysis at ZT 2.5- ZT 6.5. OKR was conducted at 0- and 30- minutes post light exposure **(Figure 6A).** As before, emixustat-treated larvae presented with a significantly reduced immediate photopic vision following dark adaptation (13.8 ± 6.1 saccades per minute, p=0.0003) which recovered 30 minutes post light exposure (23.4 ± 6.5, p<;0.0001) but not following 30 mins of removal of emixustat alone **(Figure 6B)**. Retinoid analysis revealed a reduction in 11cRAL which did not recover following a 30- minute absence of emixustat and/or 30 mins light exposure **(Figure 6C)**. A significant increase in atRP in emixustat-treated larvae in both 0- and 30-minute groups reaffirms Rpe65 inhibition (**Figure 6C**). In summary, while emixustat-treated wildtype larvae display impaired immediate photopic vision, larvae can regain vision following 30 minutes of light exposure. This light-dependent recovery of early photopic vision is not due to Rpe65 isomerase activity.

**Figure 6:**
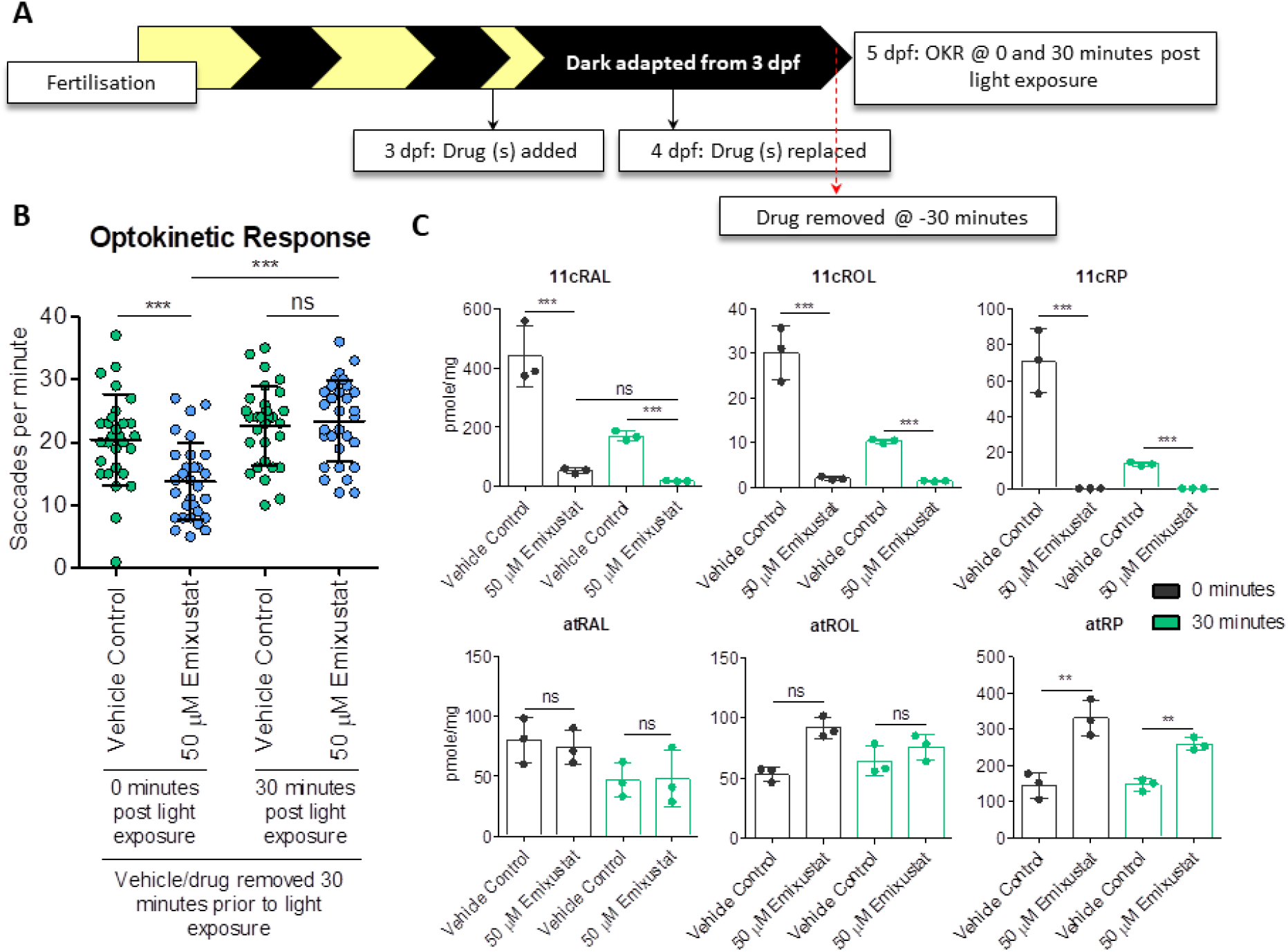
In the absence of light, 30 minutes alone is not sufficient to restore vision or 11cRAL levels. **A)** Schematic representation of experimental workflow. Zebrafish larvae are treated initially at 3 dpf and drug is replaced at 4 dpf. Larvae are incubated under dark conditions until analysis at 5 dpf. At 5 dpf, emixustat is removed under dim red light 30 minutes before larvae are subjected to OKR immediately following dark adaptation (0 minutes) or following 30 minutes light adaptation (30 minutes). **B)** Optokinetic response 30 minutes following removal of emixustat at 5 dpf larvae 0 and 30 minutes following light exposure. Data were analyzed by unpaired, two-tailed t-tests where ns=not significant, **=p<;0.01 and ***=p<0.001. N=30 larvae with 3 independent biological replicates. **C)** Retinoid profiles of 5 dpf larvae treated with 50 μM emixustat. Emixustat is removed 30 minutes before dissection at 0- or 30- minutes post light exposure. Bars represent the mean±SD of three independent experiments for each condition with 105 larval heads per biological replicate. Data were analyzed by unpaired, two-tailed t-tests where ns=not significant, **=p<0.01 and ***=p<0.001.

### Des1 is not required for cone photopic vision in zebrafish at 5 dpf

Controversially, previous studies implicated DES1 as the isomerase II in regeneration of cone visual pigments through the intraretinal visual cycle (15). To test if *degs1* is an important modulator of cone photopic vision, we created *degs1*^*−/−*^ zebrafish using CRISPR/Cas9 by engineering a 484 bp deletion in exon 2 **(Figure 7A).** Expression of *degs1* was analyzed with primers which *i)* span the deletion (primer set 2) or *ii)* nested within the deleted site (primer set 3) **(Figure 7A)**. Primer set 2 amplified a lower band of 61 bp indicating the presence of the transcript with deletion, however, it did not amplify the upper (537 bp) product observed in wildtype and *degs1*^*+/−*^ samples **(Figure 7B)**. Moreover, when primer set 3 were used, a 162 bp product is seen in wildtype and *degs1*^*+/−*^ lanes whereas no product was amplified from *degs1*^*−/−*^ samples **(Figure 7B).** No gross morphological differences were observed in *degs1*^*−/−*^ larvae at 5 dpf, compared to their unaffected siblings **(Figure 7C).** Late photopic vision at ZT 2.5 – ZT 6.5 is not significantly affected in *degs1*^*−/−*^ larvae (18.5 ± 5.4 saccades per minute) at 5 dpf compared to unaffected *degs1*^*+/+*^ or *degs1*^*+/−*^ siblings (17.1 ± 5.3 saccades per minute) **(Figure 7D-E).** We randomly drug treated offspring from a *degs1*^*+/−*^ incross with emixustat and incubated in dark **(Figure 7F).** Interestingly, neither immediate nor early photopic vision is significantly affected between the populations treated with vehicle or 50 μM emixustat **(Figure 7G).** A genotyped sample of each group validated the normal photopic response of *degs1*^*−/−*^ larvae **(Figure 7H).**

**Figure 7:**
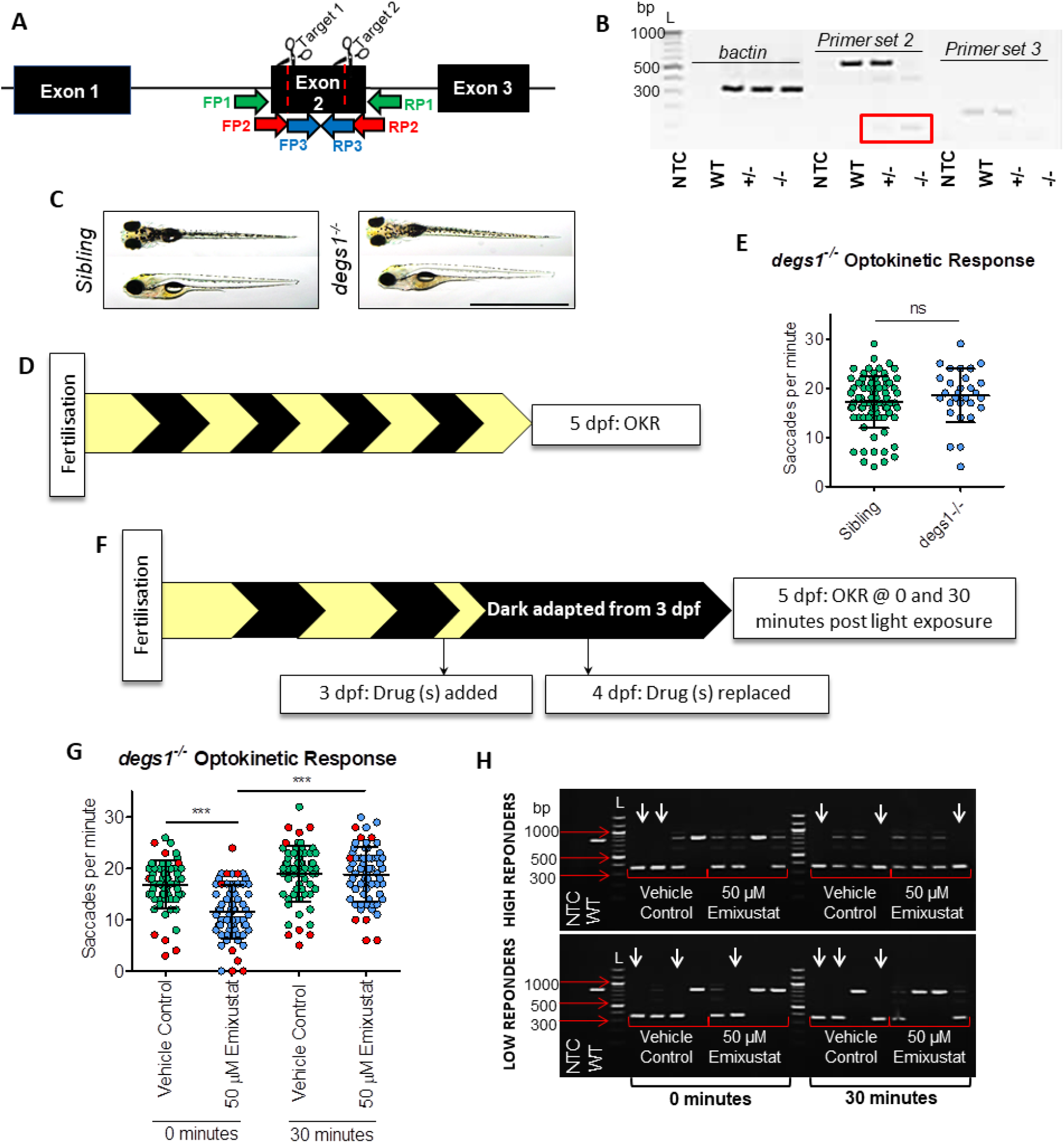
Des1 is not required for cone photopic vision in zebrafish at 5 dpf. **A)** CRISPR/Cas9 knockout strategy of *degs1* in zebrafish. Both guides were targeted to exon 2 to induce a 484 bp deletion in the *degs1* gene which contains 3 exons in total. Forward primer 1 (FP1) to reverse primer 1 (RP1) amplifies an 807 bp genomic DNA wildtype product or 323 bp product when deletion is present. For mRNA expression analysis we used forward primer 2 (FP2) to reverse primer 2 (RP2) which span the deletion and amplify a 537 bp wildtype product or 61 bp when the deletion is present. Forward primer 3 (FP3) to reverse primer 3 (RP3) is nested within the deletion and amplifies a 162 bp wildtype product. **B)** 1.5% agarose gel depicting *degs1* expression in wildtype, *degs1*^*+/−*^ or *degs^*−/−*^* zebrafish. L = 100 bp ladder. Red box highlights the 61 bp product expected following a deletion event with primer set 2 in *degs1^*+/−*^ or *degs1*^*−/−*^* samples. **C)** Dorsal and lateral brightfield microscopy images of untreated *degs1* sibling (*degs1*^*+/+*^ or *degs1*^*+/−*^) and *degs1*^*−/−*^ larvae at 5 dpf. Scale bar = 2 mm**. D)** Schematic representation of experimental workflow. *degs1*^*−/−*^ larvae are incubated under standard lighting conditions (14-hour light, 10-hour dark) until OKR analysis at 5 dpf. **E)** OKR of *degs1*^*−/−*^ and sibling larvae at 5 dpf larvae following standard 14- hour light, 10-hour dark conditions. Data were analyzed by an unpaired, a two-tailed t-test where ns= not significant (p>0.05). N≥30 larvae with 3 independent biological replicates. **F)** Schematic representation of experimental workflow. *degs1*^*−/−*^ larvae are treated initially with 50 μM emixustat at 3 dpf and drug is replaced at 4 dpf. Larvae are incubated under dark conditions until analysis at 5 dpf. **G)** At 5 dpf, a mixed population of *degs1*^*−/−*^ and unaffected sibling larvae are subjected to OKR immediately following dark adaptation (0 minutes) or following 30 minutes of light (30 minutes). Red points denote the larvae genotyped following OKR. Data were analyzed by an unpaired, a two-tailed t-test where ns=not significant (***= p<0.001). N=64 larvae per group with 2 independent biological replicates. **H)** 1.5% agarose gel highlighting genotypes of high/low responding larvae in OKR assay. White arrows denote *degs1*^*−/−*^ larvae. Eight larvae were genotyped per group (x4 ‘low responders’, x4 ‘high responders’). L=100 bp ladder.

### RPE-expressed zebrafish *rlbp1b* is not required for cone photopic vision

Co-immunoprecipitation experiments revealed that *RLBP1*-encoded CRALBP interacts with DES1 and increases the rate of 11cROL synthesis (15). Zebrafish *rlbp1* has sub-functionalized, with *rlbp1a* expressed in Muller cells and *rlbp1b* expressed in the RPE (25,26). Here, a zebrafish knockout of RPE-expressed *rlbp1b* was created using CRISPR/Cas9 which introduced a 1440 bp deletion across exon 2 and 3 **(Figure 8A).** Following *rlbp1b* knockout, *rlbp1b* gene expression was completely abolished (P<0.0001) in *rlbp1b^*−/−*^* F_2_ larval eyes. **(Figure 8C).** Gross morphology of *rlbp1b*^*−/−*^ larvae is identical to their siblings at 5 dpf **(Figure 8D).** Previous studies report a reduction in cone vision and function following *Rlbp1b* knockdown (25,26) however here, no differences in immediate (p=0.59), early (p=0.9) or late photopic vision (p=0.24) were observed when tested under standard OKR conditions **(Figure 8E-F)**.

**Figure 8:**
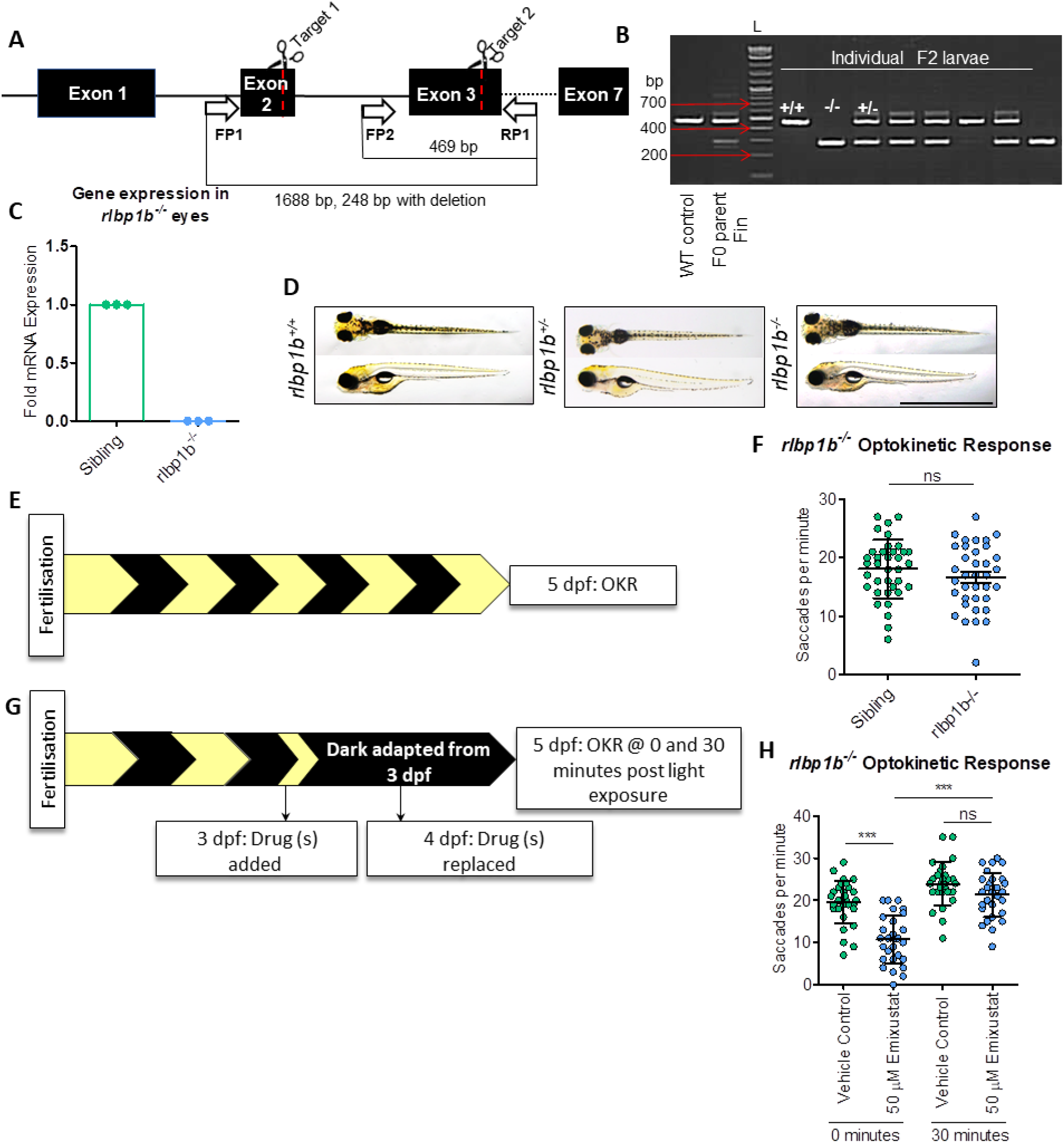
RPE-expressed *rlbp1b* is not required cone photopic vision in zebrafish at 5 dpf. **A)** CRISPR/Cas9 knockout strategy of RPE-expressed *rlbp1b*. Guides were targeted to exon 2 and exon 3 to induce a 1440 bp deletion in *rlbp1b* which contains 7 exons. Forward primer 1 (FP1) to reverse primer 1 (RP1) amplifies a 1688 bp wildtype product or 248 bp product when deletion is present. A poison forward primer (FP2) was designed in the middle of the deleted region to amplify a 469 bp wildtype product. **B)** An agarose gel (1.5%) depicting the presence of the deletion in individual F2 larval genomic DNA. A wildtype control was used to amplify the upper band (469 bp) and gDNA from the injected F0 parent fin was used as a positive control to amplify both upper and lower bands (mosaic). L = 100 bp ladder. **C)** Gene expression analysis of *rlbp1b* in *rlbp1b*^*−/−*^ eyes. Data were analyzed by unpaired, two-tailed t-tests where **=p<0.01 and ***=p<0.001. Three independent biological replicates performed with at least 10 larvae per replicate. **D)** Dorsal and lateral brightfield microscopy images of untreated wildtype, *rlbp1b*^*+/−*^ and *rlbp1b*^*−/−*^ larvae at 5 dpf. Scale bar = 2 mm. **E)** Schematic representation of experimental workflow. *rlbp1b*^*−/−*^ larvae are incubated under standard lighting conditions (14-hour light, 10-hour dark) until OKR analysis at 5 dpf. **F)** Late photopic vision as measured by OKR in 5 dpf larvae. Data were analyzed by an unpaired, a two-tailed t-test where ns=not significant (p>0.05). N=30 larvae with 3 independent biological replicates. **G)** Schematic representation of experimental workflow. *rlbp1b*^*−/−*^ larvae are treated initially with 50 μM emixustat at 3 dpf and drug is replaced at 4 dpf. Larvae are incubated under dark conditions until analysis at 5 dpf. At 5 dpf larvae are subjected to OKR immediately following dark adaptation (0 minutes) or following 30 minutes of light (30 minutes).

## Discussion

Many daily tasks are dependent on the ability of the eye to functionally adapt to changes in light. Vision in well-lit environments is known as photopic vision and is mediated by cone photoreceptors (45). Little is known regarding the biological processes underpinning functional photopic vision. Here, we apply chemical biology and genetics to dissect the contribution of light, retinoids, retinoid carrier proteins and retinoid isomerases to photopic vision *i)* immediately following dark adaptation, *ii)* after 30 minutes of light or *iii)* after ~4-6 hours of light. We exploited emixustat for its chemical ability to scavenge retinal and inhibit the RPE65 isomerase (26). Likewise, fenretinide causes depletion of vitamin A by disrupting the retinol-dependent binding of RBP4 to transthyretin (35), and also inhibits the DES1 isomerase (31–33). A1120 is a more-selective RBP4 antagonist.

### Immediate and late cone photopic vision depend on different visual cycle pathways

The ray-finned fish lineage, including zebrafish, underwent whole-genome duplication resulting in two paralogues of many teleost genes (46). In zebrafish, Rpe65a is exclusively expressed in the RPE whereas a contentious Rpe65c isoform was localized to retinal Müller glia (39). Based on this expression pattern and the requirement of mammalian cones on both the RPE and intraretinal visual cycles for complete dark adaptation (47), we predicted that emixustat could impair both immediate and possibly also early/late cone-based vision. Although emixustat inhibited cone responses immediately following exposure to light, treated larvae had normal OKR responses after prolonged light exposure. This agrees with previous genetic studies demonstrating OKR contrast sensitivity is unaffected in Rpe65a knockdown larvae raised under normal lighting conditions (20), but implicates RPE-specific Rpe65a in immediate photopic vision after dark adaptation. However, we cannot unequivocally exclude a role for zebrafish Rpe65c in Müller glia in the immediate response.

### Retinoid profiles are altered in dark-adapted zebrafish larvae treated with visual cycle isomerase inhibitors

Retinoid analysis from dark-adapted zebrafish larvae show reduced 11-*cis*-retinoids (11cROL, 11cRAL and 11cRP) with emixustat-treatment. Fenretinide also dose-dependently suppressed immediate photopic vision. However, although 11-*cis*-retinoid levels were lower in fenretinide-versus vehicle-treated larvae, the degree of suppressed vision was less than with emixustat. Levels of atROL, atRAL and atRP were also suppressed in fenretinide-versus vehicle-treated larvae, an effect not observed in emixustat-treated larvae.

### Exogenous 9cRAL supplementation restores immediate photopic vision following visual cycle inhibition

Treatment with exogenous 9cRAL recovers ERG responses in *Rpe65*^*−/−*^ mice and functional vision in *Rpe65*-deficient dogs, establishing its potential as a chromophore replacement therapy (30,47,48). In agreement, immediate photopic vision significantly improved emixustat and/or fenretinide-treated larvae supplemented with 9cRAL. Thus, the impaired cone vision we observe is a specific pharmacological effect on the visual cycle and not a result of ocular toxicity. Additionally, fenretinide-treated larvae display impaired immediate photopic vision despite relatively high 11cRAL levels. The improvement of vision with exogenous 9cRAL suggests that the 11cRAL stores present may be unavailable to cones in fenretinide-treated larvae. Improved vision in emixustat-treated larvae supplemented with 9cRAL was not coupled to increased 11cRAL levels. Instead, all 9-*cis* retinoid profiles (9cRAL, 9cROL and 9cRP) were significantly increased, indicating that the recovered vision may be mediated by iso-opsins.

### Differential requirements of Rpe65 for immediate and early cone-based photopic vision

Since emixustat did not affect late photopic vision in zebrafish, we investigated if impaired immediate photopic vision in dark-adapted animals could recover in a light-dependent, Rpe65-independent manner. Pharmacological RPE65 inhibition significantly reduces 11cRAL levels in multiple models *e.g.* by Ret-NH_2_ in zebrafish (20), by emixustat in mice (26) and by emixustat or fenretinide in *ex vivo* RPE microsomes (32). Here as expected, dark-adapted, wildtype larvae treated with emixustat display a ~10-fold reduction in 11cRAL. Interestingly, 11cRAL levels did not recover following 30 mins light exposure despite recovery of early photopic vision. A reduction in total retinoid was observed indicating that emixustat may be trapping atRAL and forming unknown or undetectable conjugates. Consistent with light overcoming emixustat-mediated visual impairment, atRAL was restored to levels similar to vehicle controls 30 minutes following light exposure, consistent with initiation of phototransduction. However, that photopic vision recovers in the light, surmounting emixustat, without an associated increase in 11cRAL levels is intriguing. Potentially, the small amount of 11cRAL available to the photopic cycle activates cone opsins in a steady state and is sufficient to sustain a stable visual response during this 30-minute light exposure. As REs support cone function (49) and are a significant source for cone pigment regeneration (50) one possibility is that 11cRP reserves were utilized upon exposure to light. However, in line with previous studies employing RPE65 inhibitors in zebrafish (49), emixustat treatment completely blocked generation of 11cRP both in the dark and light. An alternative RPE65-independent isomerization event for cone pigment regeneration was recently described in mice (18). This cone regeneration mechanisms involves Müller glia-expressed RGR-opsin, which converts atROL to 11cROL upon visible light exposure. In RPE cells, RGR opsin also affects light-dependent mobilization of atREs, suggesting it diverts substrate away from Rpe65 in RPE cells toward the Müller cell visual cycle (51).

Using genetic and pharmacological approaches, we investigate the molecular mechanisms mediating this 30-minute switch from a light-independent visual cycle, supporting immediate photopic vision, to a light-dependent visual cycle, supporting early photopic vision. When wildtype larvae were treated with the potent RBP4 antagonist A1120 alone immediate photopic vision was reduced but recovery of early photopic vision was equivalent to vehicle controls. This indicates that RBP4 is required for the non-photopic visual cycle but not for the photopic visual cycle. Furthermore, knockout of *degs1*^*−/−*^ in zebrafish showed no modulation of immediate, early or late photopic vision suggesting Des1 is not essential for either the photopic or non-photopic visual cycles. The discrepancies between Des1 pharmacological inhibition and the genetic model may be attributed to off-target effects and genetic compensation, respectively and illustrates the requirement to use these complementary approaches in parallel. Mammalian CRALBP forms a complex with DES1 (15) and is reported to regulate the cone visual cycle. Deletion of CRALBP in mice results in reduced photopic ERG responses (52). Adenovirus-mediated delivery of Müller cell CRALBP restored cone-driven responses. Here, knockout of the RPE-expressed *rlbp1b* paralogue in zebrafish did not affect immediate or early photopic vision in larvae treated with emixustat.

Significantly, fenretinide and emixustat hydrochloride have reached phase II and III clinical trials, respectively, for use in geographic atrophy. Night blindness and dry eye are side-effects of fenretinide treatment (53). Our data demonstrates that fenretinide also impairs photopic vision, thus, concerns arise over potential adverse drug reactions in the clinic with fenretinide use to treat diseases such as geographic atrophy, cancer, acne, cystic fibrosis and psoriasis. Likewise, emixustat is now in Phase III clinical trial for the inherited retinal degeneration Stargardt disease with the intention of suppressing retinoid cycling (37). We have shown that the photopic visual cycle overcomes emixustat which may have significant impact on clinical outcomes.

In summary, pharmacological and genetic models of visual cycle modulation enhanced our knowledge of the fundamental mechanisms of cone photopic vision. Differential stages of vertebrate photopic vision demand differential biochemical pathways, in light and dark. Rpe65 regenerates photopigment required by cones for immediate photopic response. In contrast, during sustained light, Rpe65 appears redundant to a light-dependent regeneration of visual pigments, a process whereby diminished 11cRAL levels are sufficient for photopic vision.

## Experimental Procedures

### Zebrafish Breeding and Maintenance

Adult zebrafish were maintained in a 14-hour light, 10-hour dark cycle in a recirculating water system at 28°C. Annual facility environmental parameters are published online: dx.doi.org/10.17504/protocols.io.3jggkjw. Wildtype (Tübingen) larvae were generated through natural spawning. Larvae were raised at 28°C in embryo medium (0.137 M NaCl, 5.4 mM KCl, 5.5 mM Na_2_HPO_4_, 0.44 mM KH_2_PO_4_, 1.3 mM CaCl_2_, 1.0 mM MgSO_4_ and 4.2 mM NaHCO_3_) containing methylene blue (Sigma Aldrich, UK).

### Ethics Statement

All experiments with zebrafish larvae were performed according to ethical exemptions granted by the UCD Animal Research Ethics Committee (AREC-Kennedy) and the Health Products Regulatory Authority (Project authorization AE18982/P062).

### Generation of mutant zebrafish

Custom crRNA guides, containing a NGG protospacer adjacent motif (PAM) were designed per gene using Benchling (www.benchling.com) CRISPR guide software. All guide sequences and PAMs are listed in *Supplementary Table 1*. The Alt-R system (Integrated DNA Technologies) was used to knockout *rlbp1b* and *degs1* in zebrafish. Briefly, 36 ng/μl crRNA and 67 ng/μl tracrRNA was complexed in nuclease-free duplex buffer by heating to 95°C for 10 minutes. Sample was cooled to room temperature and a final working concentration of 0.5 ug/μl *S. pyogenes* Cas9 nuclease (Integrated DNA technologies) was added. The mixture was heated to 37°C for 15 minutes before microinjection into wildtype embryos at one-cell stage. Injected fish were raised until adulthood and outcrossed to screen for germline transmission of the null target gene. Isogenic F_1_ heterozygous adult fish were incrossed to generate homozygous F_2_ mutants.

### Genomic DNA extraction and polymerase chain reaction

DNA was extracted from injected F0 or germline F2 rlbp1b larvae by boiling the sample at 95°C in NaOH (50 mM) (Sigma Aldrich, UK) before neutralizing with 1/10th Trizma Base (Sigma Aldrich, UK). The region of interest was amplified by polymerase chain reaction (PCR) using the following primers which span the *rlbp1b* or *degs1* mutation site: *rlbp1b* 5’-CCACAGAGTGGAACATTTCGA-3’, 5’-GATGATGCAACACTATTGCCCA-3’ and 5’-ACCTTTCAGCATTCACAATCCT-3’; *degs1* 5’-TTCTCCATGACCTTTCAGCCC-3’ and 5’-TGCTCTCACCATTGGTAGTCG-3’. Fragment sizes were run on a 1.5% agarose gel and compared against a 100 bp DNA ladder (New England Biolabs, UK).

### Total RNA extraction, cDNA synthesis and RT-PCR/qPCR

At 5 dpf, ~20-50 eyes (10-25 larvae) were enucleated and stored in RNAlater (Qiagen, Germany) individually at 4°C. Larval trunks were genotyped as previously described. Eyes were pooled and total RNA was extracted using mirVana™ miRNA Isolation Kit (ThermoFisher Scientific) as per the manufacturer’s instructions. NaOAc (3M) and 100% ethanol were added to the eluted RNA, vortexed and left overnight at −20°C. Samples were centrifuged at 14000 rpm for 60 minutes at 4°C, supernatant removed before the pellet was resuspended in 80% ethanol and centrifugation for a further 60 minutes at 14000 rpm. Supernatant was discarded and the pellet was resuspended in ultrapure water. Total RNA concentration was quantified at 260 nm (Spectrophotometer ND- 1000) and samples stored at −80°C until further use. cDNA was synthesized using PrimeScript™ RT reagent Kit (Perfect Real Time) (TAKARA, Japan). QRT-PCR reactions were made up on ice with 0.2 μl forward and reverse primer: *rlbp1b* 5’-TGAGCTTGCTAAAGGTGTTCAGG-3’ and 5’-TCAGGATAATCCCGTCTGAAGC-3’, 5 μl Sybr Green Master Mix (Thermo Fisher Scientific, Massachusetts, USA), 3.6 μl RNAse- free water and 1 μl cDNA template (12.5 ng) was added per sample. QRT-PCR cycles were carried out using the following parameters: 50°C for 2 min, 95°C for 10 min, 95°C for 15 s with 40 repeats and 60°C for 1 min on a Quant Studio 7 Flex Real-Time PCR System. To evaluate *degs1* expression, the cDNA was amplified by PCR using the following primer sets: *degs1 forward 2:* 5’-GTCAGCACGATGGTGGT GTC-3’*, degs1 reverse 2:* 5’-TGGAGACCCATGCCCAGCAT-3’*; degs1 forward 3:* 5’-GGACGTGGACATCCCCACTGA −3’, *degs1 reverse 3:* 5’-AGCTGGATGGCCACGTTCAG −3’. Samples were run on a 1.5% agarose gel and compared against a 100 bp DNA ladder (New England Biolabs, UK).

### Drug Preparation and Treatment

Wildtype, *rlbp1b*^*−/−*^ and *degs1*^*−/−*^ larvae were raised from embryos under standard 14-hour light, 10- hour dark conditions. At 3 dpf, five larvae per well were placed in 48 well cell culture plates (Greiner Bio-one, Austria). Emixustat hydrochloride (Medchem, New Jersey, USA), fenretinide (Cayman Chemical, Michigan, USA) and A1120 (Sigma Aldrich, UK) were prepared in embryo medium and 400 μl drug solution at 1-50 μM in 0.01%- 0.5% DMSO was added to the wells. For emixustat and fenretinide combinations, 1-10 μM fenretinide was added 15 mins prior to the addition of 50 μM emixustat. Larvae were initially treated at 74-76 hpf and the drug was replaced at 98-100 hpf. 10 μM 9-*cis*-retinal (Sigma Aldrich, UK) was added to embryo medium four hours following primary drug treatment (*i.e.* 78-80 hpf and 102-104 hpf). 9- *cis*-retinal was handled under dim red light.

### Visual Behavior Analysis

To measure optokinetic response (OKR), single larvae were immobilized in pre-warmed 9% methylcellulose (Sigma Aldrich, UK) and placed in the center of a rotating drum containing vertical 1 cm thick black and white stripes. The angle subtended was 0.58 rad (33.7°). The drum was rotated at 18 rpm for 30 seconds in a clockwise and 30 seconds in a counterclockwise direction. Saccadic eye movements per minute were recorded manually. All OKR measurements were performed in larvae at 5 dpf between 10 am (ZT 2.5) and 2 pm (ZT 6.5).

### Histological Analysis

Larvae were fixed in glass vials containing 2.5% glutaraldehyde, 2% paraformaldehyde and 0.1% Sorenson’s phosphate buffer (pH 7.3) and were placed at 4°C. Samples were washed before post-fixation in 1% osmium tetroxide (Sigma Aldrich, UK) and dehydrated using an ethanol gradient (30%, 50%, 70%, 90%, 100%). Samples were transferred to propylene oxide (Sigma Aldrich, UK), embedded in agar epoxy resin overnight and 0.5 μm sections were cut using an ultramicrotome (Leica EM UC6). Sections were placed on glass slides and stained with toluidine blue (Sigma Aldrich, UK) for 30-60 seconds and imaged using a Leica DMLB bright field illumination microscope with a Leica DFC 480 camera attachment.

For transmission electron microscopy (TEM), 0.08 μm sections were cut using a diamond knife and a Leica EM UC6 ultramicrotome. Sections were transferred to a support grid, stained with uranyl acetate and lead citrate, and imaged on a FEI-Tecnai 12 BioTwin transmission electron microscope (FEI Electron Optics).

Drug treated larvae were fixed at 5 dpf with 4% paraformaldehyde (PFA; Sigma Aldrich, UK). Larvae were cryoprotected in a sucrose series before embedding in optimal cutting temperature (OCT) medium (VWR, Pennsylvania, United States). Sections were cut to 14 μm and thaw-mounted onto charged superfrost plus slides (Fisher Scientific, Pittsburgh, USA). Sections were rehydrated using 1X phosphate buffered saline supplemented with tween 20 (Sigma Aldrich, UK) (0.01% PBST) before blocking in 10% goat serum (Sigma Aldrich, UK) for 1 hour. Monoclonal zpr-1 was used at 1:200 (Zebrafish International Resource Center, Eugene, USA). Slides were washed before incubating with the secondary antibody (Alexa Fluor 488 anti-mouse IgG). Primary and secondary antibodies were diluted in blocking solution. Slides were mounted using Vectashield; (Vector Laboratories, California, USA) before imaging using a Zeiss LSM510 confocal microscope (Carl Zeiss AG, Germany) with a 63X objective.

### Retinoid Quantification

For retinoid analysis, 105 larvae per treatment condition at 5 dpf were euthanized in ice water before their heads were collected under dim red light. For normal-phase HPLC analysis of retinoids, all retinoid extractions were carried out under dim red light in a dark room. Zebrafish heads were stored at −80 °C, thawed on ice, and homogenized in 500 μl of phosphate buffered saline (PBS) using a glass tissue grinder (Kontes). Protein concentration was determinated using 50 μl of sample and a Micro BCA Protein Assay Kit (Pierce, US). To the remaining 450 μL of homogenate 25 μL of 5% SDS (~0.2% SDS final concentration) and 50 μL of brine were added and samples briefly mixed. Hydroxylamine hydrochloride (500 μL of 1.0 M solution in pH 7.0 PBS) was added to generate retinal oximes and samples were vortexed and incubated at room temperature for 15 minutes. The aqueous phase was quenched and diluted using 2 ml of cold methanol. The samples were twice extracted by addition of 2 ml aliquots of hexane, brief vortexing, and centrifugation at 3000 × g for five minutes to separate the phases. Pooled hexane extracts were added to 13 × 100 mm borosilicate test tubes and evaporated to dryness under a stream of nitrogen. Dried samples were dissolved in 100 μl hexane and analyzed by normal-phase HPLC using an Agilent 1100 series chromatograph equipped with a Supelcosil LC-Si column (4.6 × 250 mm, 5 μm) using a 0.2–10% dioxane gradient in hexane at a flow rate of two mL per minute. The eluted mobile phase was analyzed using a photodiode-array detector. Spectra (210–450 nm) were acquired for all eluted peaks. The identity of eluted peaks was established by comparison to spectra and elution times of known authentic retinoid standards. Retinoid amounts were quantitated by comparing their respective peak areas to calibration curves established with retinoid standards.

### Photostability of Emixustat Hydrochloride

Liquid chromatography used a Dionex Ultimate 3000 RSLC micro-LC system (Thermo Fisher Scientific, US). Separation was performed using a Thermo Acclaim RSLC 120 2.2 μm, 120 A, 1.0 × 100 mm LC column (Thermo Fisher Scientific, US). LC parameters were: elution mode: gradient (from 90% A to 90% B in 15 min), flow rate: 45 μL/min, column temperature: 3°C, detector wavelength: 230 nm and injection volume: 2 μL. Mobile phases were: A: H_2_O + 0.1% formic acid (FA) (v/v) B: Acetonitrile + 0.1% FA (v/v). Mass Spectrometry analysis was performed using a LTQ-XL-Orbital-XL (Thermo Fisher Scientific, US) equipped with HESI II ESI ion source. Source parameters were: Ion spray voltage (V): 4.6, capillary Temp (°C): 280, sheath gas flow gas flow (Arb): 20 and auxiliary gas flow (Arb): 8. Resolution was set to 30,000 in FT mode and mass range set to 150-500 Da in positive ion mode. Peak area of emixustat protonated mass was calculated and recorded using Xcalibur software (version 2.21). The protonated mass ions were identified using an error tolerance of 5 ppm for their corresponding theoretical monoisotopic mass.

For Nuclear Magnetic Resonance (NMR) analysis, emixustat (7 mg) was dissolved in DMSO-d6 (600 μl) under dim red light and covered with aluminum foil until ^1^H NMR and HSQC experiments were performed. The NMR tube was exposed to visible light for 30 minutes, followed by a repeat NMR analysis. ^1^ H NMR spectra were measured in the solvent stated at 400 MHz. Chemical shifts (δ) are quoted in parts per million (ppm) referenced to residual solvent peak (e.g., DMSO-d^6^: ^1^H – 2.50 ppm and 3.33 ppm).

### Statistical Analysis

Statistical analysis was performed using GraphPad Prism™ software (GraphPad, San Diego, CA). A two-tailed student’s unpaired t-tsest was applied comparing two experimental groups. For analysis involving more than two independent groups, a one-way ANOVA and Dunnett’s multiple comparison test or Bonferroni post hoc test was performed. Experiments comprised of two variables were tested for significance using a two-way ANOVA and a Bonferroni post hoc test. All data are presented as mean ± standard deviation (SD). Statistical significance was assigned to p-values of p <; 0.05 = *, p < 0.01 = ** or p < 0.001 = ***.

## Acknowledgements

We thank the UCD Biomedical Facility technicians for zebrafish maintenance and the UCD Conway Institute Imaging Core, with particular thanks to Dr. Dimitri Scholz and Dr. Niamh Stephens. We thank Prof. Jack Saari for helpful discussions surrounding this project and Dr. Ross Collery for proof-reading the manuscript. This project received funding from an Irish Research Council/Fighting Blindness grant (EPSPG/2017/276), the European Union’s Horizon 2020 Research and Innovation Staff Exchange programme under grant agreement No. 734907 (RISE/3D-NEONET project), and by NEI, National Institutes of Health Grants R01-EY024379 (to G.H.T.) NEI, National Institutes of Health Core Grant P30-EY000331; and Research to Prevent Blindness unrestricted grant (to the Jules Stein Eye Institute). G.H.T. is the Charles Kenneth Feldman Professor of Ophthalmology at UCLA.

## Conflict of interest

The authors declare that they have no conflicts of interest with the contents of this article

## Abbreviations

AMD: Age-related macular degeneration
atROL: all-*trans*-retinol
A2E: *N*-retinylidene-*N*-retinylethanolamine
atRE: all-trans-retinyl esters
atRP: all-trans-retinyl palmitate
CRISPR/Cas9: Clustered regularly interspaced short palindromic repeats/ CRISPR-associated protein 9
DES1: Dihydroceramide desaturase-1
LRAT: Lecithin retinol acyltransferase
MFAT: Multifunctional O-acyltransferase
OKR: Optokinetic Response
RBP4: Retinol-binding protein 4
RPE: Retinal Pigment Epithelium
TTR: Transthyretin
RBP4: Retinol-binding protein 4
ZT: Zeitgeber
9cRAL: 9-cis-retinal
11cRAL: 11-cis-retinal
11cROL: 11-cis-retinol
11cRE: 11-cis-retinyl ester
11cRP: 11-cis-retinyl palmitate
13cRAL: 13-cis-retinal

