## Supplementary Figure 1 for "Non-photopic and photopic visual cycles differentially regulate immediate, early and late-phases of cone photoreceptor-mediated vision"

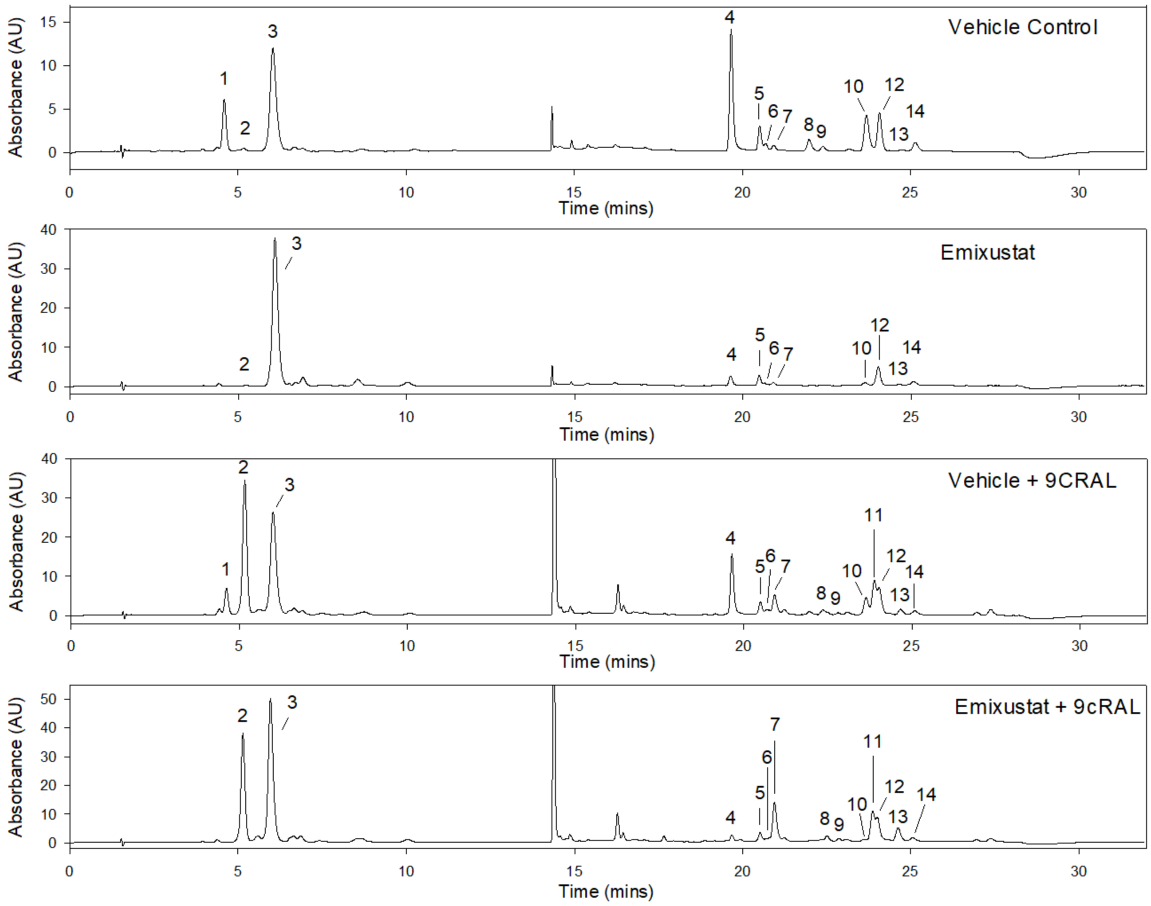


**Supplementary Figure 1:** HPLC chromatograms of retinoids present in 5 dpf zebrafish heads following treatment with emixustat + exogenous 9cRAL. Identification of peaks 1-14 as follows: 1) 11cRP, 2) 9cRP, 3) atRP, 4) syn-11cROX, 5) syn-atROX, 6) syn-13cROX, 7) syn-9cROX, 8) 11cROL, 9) anti-13cROX, 10) anti-11cROX, 11) 9cROL, 12) atROL, 13) anti-9cROX, 14) anti-atROX.
