## Supplementary Table 1 for "Non-photopic and photopic visual cycles differentially regulate immediate, early and late-phases of cone photoreceptor-mediated vision"

| Gene Symbol | Strand | Sequence (5'-3') | PAM |
|---|---|---|---|
| ***degs1*** | + | *ACCTGGTGAAGGATCTGTCC* | TGG |
| | - | *CCCATCATCATGTAGACCAC* | AGG |
| ***rlbp1b*** | - | *AGGCTTCAAACACGGGCCCA* | TGG |
| | + | *ACTGGAGATGAGCTTGCTAA* | AGG |

**Supplementary Table 1:** List of CRISPR guide sequences for *degs1* and *rlbp1b* knockout zebrafish.
